# Fungicide drives *de novo* evolution of multidrug resistance in the plant growth promoting rhizobacterium, *Pseudomonas fluorescens*

**DOI:** 10.64898/2026.04.01.715857

**Authors:** Matthew Kelbrick, James P. J. Hall, Siobhán O’Brien

## Abstract

Plant growth-promoting rhizobacteria (PGPR) are key members of soil microbial communities, supporting nutrient cycling, plant health, and productivity. In agricultural soils, these beneficial bacteria are often exposed to multiple stressors simultaneously, including fungicides, antibiotics, and rising temperatures. Despite their ecological importance, little is known about how PGPR respond evolutionarily to such combined stressors. Here, we investigated how the model PGPR *Pseudomonas fluorescens* SBW25 evolves resistance to the fungicide formulation Fubol Gold (metalaxyl-M + mancozeb), under ambient or warming soil conditions. Using a 16-week experimental evolution in soil microcosms with a fully factorial design (fungicide ± warming), we assessed the evolution of fungicide resistance via phenotypic assays and whole-genome sequencing. Fungicide exposure rapidly selected for increased fungicide resistance, detectable as early as week 4, and co-selected for multidrug resistance, likely through mutations in a *mexS* ortholog that cause efflux pump overexpression. Warming did not alter the evolution of fungicide resistance; however, populations subjected to both fungicide and warming stress went extinct more rapidly, indicating that population evolutionary rescue was less effective under dual stress. Our findings show that fungicides alone can drive multidrug resistance in beneficial soil bacteria, suggesting that strategies to tackle AMR in agriculture should also consider non-antibiotic drivers of resistance.

## 1. Introduction

Fungicides, including azoles, dithiocarbamates and phenylamides, are widely applied in agricultural systems to control plant-pathogenic fungi and safeguard crop yields [1, 2]. Although designed to target fungal pathogens, fungicides inevitably enter soil environments where they interact with diverse non-target microorganisms [2]. Soil bacterial communities play essential roles in nutrient cycling, organic matter turnover and plant health [3], and there is increasing evidence that fungicide application can alter soil microbiome structure and function. Experimental and field studies show that fungicides shift bacterial community composition, reduce alpha diversity and disrupt microbial interaction networks in agricultural soils [4-7]. Plant growth-promoting rhizobacteria (PGPR), are a functionally important group of soil bacteria that enhance plant productivity through nutrient mobilisation, phytohormone production and pathogen suppression [8]. Fungicide resistance has been identified in several PGPR taxa, raising the possibility that resistant strains could buffer crops against the negative side effects of chemical disease control [9, 10]. However, the ecological and evolutionary dynamics underpinning fungicide resistance in PGPR, especially under future climate conditions, remain poorly understood.

Agricultural soils are inherently multistressor environments in which microorganisms are exposed not only to pesticides but also to fertiliser inputs, heavy metals and rising temperatures associated with climate change [11]. Adaptation under multiple stressors can be constrained by trade-offs, such as antagonistic pleiotropy, where mutations that confer resistance to one stressor reduce tolerance to another [12]. Genetic correlations, epistasis, and population bottlenecks may further limit adaptive potential [13]. Conversely, resistance to multiple stressors may arise through shared physiological mechanisms, such as the upregulation of multidrug efflux pumps [14-16] or broad-spectrum hydrolases capable of degrading diverse xenobiotics [17]. Co-regulation among stress response pathways may also accelerate adaptation when exposure to one stressor primes tolerance to another, as has been observed for elevated temperature and certain antibiotic classes [18]. Despite these contrasting possibilities, we lack empirical tests of how warming shapes fungicide resistance evolution in beneficial soil bacteria.

Here, we investigate whether warming temperatures constrain fungicide resistance evolution in the model PGPR *Pseudomonas fluorescens* using experimental evolution in compost soil microcosms [19, 20]. Our microcosm system mimics the spatial structure and resource heterogeneity of natural soils while maintaining high replication and experimental control under laboratory conditions. We first identify a fungicide formulation that has a strong inhibitory effect on *P. fluorescens* (Fubol Gold – a commercially used fungicide formulation containing the phenylamide metalaxyl-M and the dithiocarbamate mancozeb, applied to control oomycete pathogens such as potato late blight). Next, we used experimental evolution to test whether warming temperatures constrain fungicide resistance, using a combination of phenotypic assays and whole genome resequencing on evolved and ancestral PGPR. Finally, we test whether fungicide-evolved isolates display cross-resistance to a panel of antibiotics. Our findings show that fungicide drives selection for both fungicide and antibiotic resistance in *P. fluorescens*, irrespective of warming.

## 2. Methods

### 2.1: Bacterial strains

For experimental evolution, we used *P. fluorescens* SBW25 carrying a gentamicin-resistance cassette (SBW25::Gm^R^) [21]. Because the ancestral strain showed poor survival in soil, we instead used a SBW25::Gm^R^ isolate that had previously adapted to John Innes No. 2 soil-based compost. This isolate, A.21.65.G.003, was retrieved from a plasmid-free population of *P. fluorescens* SBW25 that had evolved alongside *P. putida* KT2440 in control soil microcosms as described in [19, 22]. Previous whole-genome sequencing (Short Read Archive accession number ERR1554495) identified three nonsynonymous mutations in this soil-adpated SBW25::Gm^R^ relative to the reference SBW25 genome: Phe133Val in PFLU_5421 (putative peptidoglycan-related gene), Asp225Ala in PFLU_6004 (conserved hypothetical), and Thr128Pro in PFLU_0941 (putative peptidoglycan glycosyltransferase) [20], which were also detected in all eight ancestral clones in our evolution experiment (see section 2.2, Supplementary data S3). In preliminary assays, the soil-adapted SBW25::Gm^R^ was recoverable in all replicates (n = 12).

### 2.2: Quantifying inhibitory effects of four different fungicide formulations on bacterial growth

We assessed the effects of four fungicides used in the EU or UK at the time of this study: *Amistar* (Syngenta; azoxystrobin), *Fubol Gold* (Syngenta; mancozeb, metalaxyl-M), *ProPlant* (Dejex; propamocarb hydrochloride), and *Subdue* (Dejex; metalaxyl-M) on *P. fluorescens* final densities in soil microcosms. 10 g of John Innes No. 2 soil (J. Arthur Bowers) was placed in 30 ml glass vials, autoclaved twice at 126 °C for 15 min, and left at room temperature for five days to allow reoxidation of heat-generated toxic compounds [23]. Fungicides were mixed with sterile distilled water at a 100:1 (w/w) ratio. Each soil microcosm received 900 µl fungicide solution, giving a final concentration of 0.1% (w/w) in soil. Control microcosms received 900 µl sterile water. Three replicate microcosms were prepared per treatment (15 microcosms total), vortexed for 30 s, and equilibrated for 1 h.

A frozen stock of *P. fluorescens* was streaked onto Lysogeny Broth Agar (LBA) and incubated at 26 °C for 48 h. A single colony was cultured in 10 ml Lysogeny Broth (LB) at 26 °C, 180 rpm for 24 h, centrifuged (3000 × g, 10 min), washed twice in M9 buffer, and resuspended in 10 ml M9. Each soil microcosm was inoculated with 100 µl of this culture (10□ colony-froming units (CFUs) ml^−1^) so the starting density of *P. fluorescens* was 10^3^ CFUs g^−1^ soil. Microcosms were vortexed for 30 s, and incubated at 26 °C and 70% humidity for four days. After incubation, 20 sterile glass beads (5 mm; Witeg) and 10 ml M9 buffer were added, vortexed for 45s, and left for 30 min to facilitate the separation of the supernatant from the soil wash. The supernatant was serially diluted and plated (50 µl) on LBA containing 25 µg ml^−1^ gentamicin (Melford) to ensure selective growth of our *P. fluorescens* strain. Plates were incubated for 48 h at 26 °C, and CFU’s g^−1^ soil were enumerated. All fungicide formulations significantly decreased *P. fluorescens* final densities compared to fungicide-free controls (Fubol Gold; t-test: t = 143.73, p.adj < 0.001; Amistar; t-test: t = 28.955, p.adj < 0.001, Subdue; t = 33.379, p.adj < 0.001, ProPlant; t-test: t = 20.645, p.adj < 0.001, Figure S1). For our remaining experiment, we chose to focus on Fubol Gold, as it had the strongest inhibitory effect on *P. fluorescens*.

### 2.3: Experimental evolution of P. fluorescens in soil

*P. fluorescens* was experimentally evolved in sterile soil microcosms under four treatments: control, fungicide, warming, and fungicide + warming, using a fully factorial design (8 replicates per treatment, 32 microcosms in total). Fungicide concentrations were increased every 4 weeks: (week 1-4: 10 mg kg−1 (0.001%); week 4-8: 50 mg kg−1 (0.005%); week 8-12: 100 mg kg−1 (0.01%); week 12-16: 500 mg kg−1 (0.05%) (w/w). Preliminary experiments showed a ∼6000-fold reduction in *P. fluorescens* densities at 0.1% Fubol Gold; therefore, concentrations were kept below this level during the evolution experiment to prevent extinction. Warming-treated microcosms increased from 26 °C in increments of 0.5 °C every seven days, until a maximum temperature of 34 °C at week 16.

A 1 g mL^−1^ Fubol Gold stock was prepared in sterile distilled water and diluted to the desired concentration, starting at 10 mg kg^−1^ soil (0.001% w/w) at week 1. Sixteen 10 g sterile soil microcosms received 900 µL of fungicide solution, and controls received sterile water. Microcosms were vortexed for 30 s and equilibrated for 1 h before inoculation.

A frozen stock of *P. fluorescens* was streaked on LBA supplemented with 25 µg ml^−1^ gentamicin and incubated for 48 h at 26 °C. Eight colonies were randomly selected and cultured individually in 10 ml LB at 26 °C with shaking (180 rpm) for 24 h, centrifuged (3000 × g, 10 min), washed twice with M9 buffer, and resuspended in 10 ml M9. 100 µl (10^5^ CFU ml^−1^) of each clone was inoculated into four soil microcosms each (one soil microcosm per treatment) so final bacterial density was 10^3^ g^−1^ soil. Microcosms were vortexed for 30 s and incubated at 26 °C and 70% relative humidity.

Populations were passaged every 7 days for 16 weeks (112 days). 10ml M9 buffer and 20 sterile glass beads were added to each microcosm, vortexed for 45 s, and allowed to settle for 30 min. A 100 µL aliquot of the soil wash was transferred to fresh microcosms with the appropriate fungicide concentration and vortexed for 30 s. Every 4 weeks, 150 µL of soil wash was cryopreserved in 20% (w/v) glycerol at −80 °C. Population densities (CFU g^−1^ soil) were quantified every 4 weeks by serial dilution of 20 µL soil wash in M9, plating on gentamicin-supplemented LBA, and incubating for 24 h. To verify identity of our inoculated SBW25::Gm^R^ strain, 12 clones (3 per treatment) were randomly selected and confirmed as SBW25::Gm^R^ by PCR targeting the mini-Tn7 GentR cassette (Supplementary Methods).

### 2.4: Genome resequencing evolved and ancestral P. fluorescens clones

To identify treatment-specific mutational patterns, 2 colonies were randomly selected from each population at the final time point (week 16); for populations that went extinct earlier, clones were sampled from the last viable time point. All eight ancestral clones were also sequenced. Frozen population stocks were thawed, serially diluted, plated on LBA, and incubated at 26 °C for 48 h. Two colonies per plate were cultured in 5 mL LB (26 °C, 180 rpm, 24 h) and cryopreserved (1 mL culture mixed with 500 µL of 60% w/v glycerol, −80 °C). Clones were re-streaked on LBA, incubated for 48 h, and biomass was harvested into 500 µL DNA/RNA Shield (MicrobesNG). Genomic DNA was extracted following MicrobesNG protocols and sequenced on an Illumina NovaSeq 6000 platform (2 × 250 bp; >30× coverage). Reads were trimmed with Trimmomatic v0.30 and quality-checked using FastQC. Variants were called with breseq v0.37.1 [24] against the *P. fluorescens* SBW25 reference genome (AM181176). Two contaminant (non-Pseudomonas) clones (both from warming population 4) were excluded from analysis. All reads have been uploaded to the NCBI SRA database (BioProject ID: XXXXXX).

### 2.5: Quantifying fungicide minimium inhibitory concentrations for evolved and ancestral clones

To test whether (i) clones from fungicide-evolved treatments differed in fungicide resistance versus control-evolved clones and (ii) whether fungicide-evolved clones carrying mutations in *PFLU_3160* exhibited increased fungicide resistance compared to wildtype, we performed a minimal fungicide inhibitory concentration (MIC) assay. One clone was selected at random from each fungicide-evolved, fungicide+warming evolved, control-evolved and ancestral populations (32 clones in total, 8 clones per treatment). These clones also varied with respect to *PFLU_3160* mutations (13/16 clones had *PFLU_3160* mutation in fungicide treatments, while all ancestral and control clones had wildtype *PFLU_3160*). Clones were streaked from frozen stocks onto LBA plates and incubated for 48 h at 26 °C. Several colonies from each plate were collected using sterile cotton swabs and suspended in 10 ml 0.85% (w/v) saline to achieve a cell density equivalent to a McFarland 0.5 standard.

A 65.5 mg ml^−1^ solution of fungicide was prepared in × 2 LB broth (diluted to × 1 when culture in saline was added). The fungicide solution was serially diluted in a 96-well plate, with each dilution transferring 100 µl fungicide solution into 100 µl LB (for a 2-fold dilution series); this was repeated 24 times until fungicide concentration was reduced to 0.015625 µg ml^−1^. A 100µl sample was removed from the final well and discarded. This dilution series was replicated four times, three as experimental replicates and one as an uninoculated control. A 100 µl sample of standardised bacterial culture in 0.85% saline (or 0.85% saline for controls) was added to each of the wells (diluting LB to × 1). Thus, the final fungicide concentration in wells was between 32.25 mg ml^−1^ and 0.0078 µg ml^−1^ in two-fold increments. Plates were incubated at 26 °C and 600RPM for 24 h using a microplate shaker (Stuart) at 600 RPM. Growth inhibition was visually assessed, with the lowest fungicide concentration showing no visible growth (clear well) recorded as the minimal inhibitory concentration (MIC) in µg ml^−1^ for each clone.

### 2.6: Quantifying antibiotic resistance for evolved and ancestral clones

We tested whether (i) fungicide evolved clones had increased antimicrobial resistance compared to control-evolved clones, and (ii) whether *PFLU_3160* mutants had increased resistance compared to wildtype, using antimicrobial resistance (AMR) disk diffusion assays in accordance with EUCAST guidelines [25]. Clones were standardised to a cell density equivalent to a McFarland 0.5 standard and using a sterile cotton swab, cultures were spread onto square agar plates (50 ml LBA) in three directions to ensure complete coverage. Then, either MAST antibiotic rings (impregnated with Colistin Sulphate 25 µg, Streptomycin 10 µg, Sulphatriad 200 µg or Tetracycline 25 µg) or disks (impregnated with Nalidixic acid 30 µg, or Chloramphenicol 30 µg) were firmly placed onto the agar surface using autoclaved sterile tweezers. Each disk assay was repeated three times per clone. Plates were incubated for 24 h at 26 °C. After 24 h, inhibition halos were measured to the nearest millimetre using a ruler.

### 2.7: Statistical analysis

#### (i) Fungicide inhibition tests

To assess the effects of four different fungicide formulations on *P. fluorescens* growth, we conducted pairwise *t*-tests on log_10_-transformed CFU g^−1^ soil data, comparing each formulation with the fungicide-free control. *P*-values were adjusted for multiple comparisons using Bonferroni correction.

#### (ii) Population densities over time

Population extinction dynamics were analysed using survival analysis, with extinction treated as the event and surviving populations right-censored at the final sampling point. Extinction was defined as the absence of viable cells by plating. Kaplan–Meier survival curves were estimated using survfit() function in the survival package [26] and visualised using ggsurvplot() in the survminer package [27]. Differences in survival among warming and fungicide treatments were assessed using log-rank tests (survdiff() function in the survival package) which account for both the frequency and timing of extinction events. The pairwise_survdiff() function from the survminer package was used to test for pairwise comparisons between treatments, with Bonferroni correction for multiple testing.

#### (iii) Genome resequencing

To accommodate mutations in our soil-adapted SBW25 ancestor relative to the reference genome, and to control for spurious prediction of mutations in repetitive or difficult-to-map parts of the reference, any mutations identified in our 8 ancestral clones relative to the SBW25 reference sequence, including mutations with ‘marginal’ or ‘unassigned’ evidence, were pooled and removed from all clones in the evolved variant dataset prior to statistical analysis. To detect if mutational patterns among evolved clones were treatment specific we used a factorial PERMANOVA with 10,000 permutations using the ‘adonis’ function from the ‘vegan’ v 2.6-4 package in R [28]. Significance of individual model terms was assessed using marginal tests (by = “terms”). Homogeneity of multivariate dispersion among treatment groups was evaluated using the betadisper function, followed by ANOVA, to confirm that significant PERMANOVA results were not driven by differences in within-group variance.

To identify putative treatment-specific parallel mutations, we identified candidate mutations as those which appeared in >1 population within a treatment and 0 in the alternative treatment. Directly adjacent genes belonging to the same operon with overlapping functions (e.g. *actP* and *PFLU_1813*) were grouped and treated as a single mutational target. Candidates were tested using Fisher’s exact test, with p-values corrected for multiple testing using a Bonferroni-adjustment. In our dataset of 7 parallel targets, two targets consisted of large deletions in intergenic regions including (i) a putative 133bp deletion in the intergenic region *PFLU_2441-PFLU_2442* (downstream of two LysR family transcriptional regulators), detected in 32/62 clones across all 4 treatments, and (ii) a putative 167 – 249bp deletion in intergenic region *PFLU_4254-PFLU_4255* (a transposase and an integral membrane protein coding gene respectively; 11/62 clones across all 4 treatments). Notably, these deletions were predicted in repeat-rich regions prone to spurious mutation calls [24, 29] so were removed from our final dataset of parallel mutational targets.

#### (iv) Resistance assays

Mode fungicide MIC (3 relicates per clone) was taken as the MIC for each clone. We tested for differences in log_2_ fungicide MIC between evolution treatments (control, fungicide and fungicide+warming) using ANOVA and Tukey tests, confirming normality of residuals using QQ-plots and Shapiro-Wilk normality tests (W=0.95). Differences in log_2_ MIC between *PFLU_3160* mutants and wildtype clones was tested using a Welch two sample t-test, to allow for unequal variances between treatments. To test the relative contributions of evolutionary treatment and mutation status on fungicide MIC, we used a linear model (log_2_ MIC ∼ treatment + mutation status) and likelihood ratio tests.

We tested whether antimicrobial resistance (quantified as the mean zone of inhibition in millimeters) differed between treatments (control, fungicide, and fungicide + warming) using Kruskal-Wallis with Bonferonni correction, and Dunn posthoc tests. All clones (including ancestral strains) were completely resistant to ampicillin, cephalothin, and cotrimoxazole (zone of inhibition = 0 mm for all clones); these antibiotics were therefore excluded from statistical analyses. Differences in resistance between *PFLU_3160* mutants and wildtype clones was tested for each antibiotic using Wilcoxin Rank-Sum tests with Bonferonni correction.

## 3. Results

### 3.1 Dual stressors accelerate population extinction

Treatment (fungicide, warming, dual stressors or stress-free control) significantly affected extinction risk in our evolving populations (log-rank test: χ^2^_3_ = 31.5, p < 0.0001, Figure S2). Populations in the control treatment remained fully viable throughout the experiment, with densities consistently high, ranging from 3.0 × 10 □ to 9.0 × 10 □ CFU g^−1^ soil (Figure S3). All single-stressor populations remained viable until week 16, when extinction occurred in all fungicide populations and 4/8 warming populations. In contrast, dual-stressor populations had a median survival of 12 weeks, with 2/8 populations reaching extinction by week 8. When stressors were imposed independently, fungicide increased extinction risk relative to controls (p<0.001), but warming did not (p>0.05). Dual stressor treatments had increased extinction risk relative to controls (p<0.001), fungicide only (p<0.05) and warming only (p<0.01) (Figure S2). Our data shows that extinction rates in dual-stressor populations could not have been predicted from single-stressor dynamics alone.

### 3.2 Pinpointing genetic responses to selection in a multi-stressor environment

We hypothesised that, prioir to extinction, populations could evolve by acquiring mutations to adapt to the increasing stress. To identify treatment-specific patterns of mutation, we selected two clones from each evolved population at the final time point prior to extinction for whole-genome resequencing. Across evolved clones, we detected 152 mutations absent in ancestral clones, comprising gene deletions (53%), insertions (11%) and single nucleotide polymorphisms (37%), including non-synonymous (45%), synonymous (16%), intergenic (9%) and nonsense (30%) (Supplementary data S1, Figure S4). Evolved clones had between 0 and 6 mutations with a median of 2 mutations per clone. We did not detect any mutations in one clone (B8Y; fungicide-only treatment).

We identified mutations in 36 distinct loci across all evolved clones, with 7 targets mutated in more than one population (i.e. parallel evolution) (Figure 1, Supplementary data S2). Genome-wide patterns of mutation were significantly associated with fungicide treatment (permutational MANOVA: effect of fungicide, R^2^=26.8,F_1,27_ = 10.54, p < 0.0001), irrespective of warming (fungicide × warming, R^2^=0.02, F_1,27_ = 0.96, p > 0.05). We could not detect any statistical association between warming and the pattern of mutations (effect of warming, R^2^=0.02, F_1,27_ = 0.83, p > 0.05).

**Figure 1:**
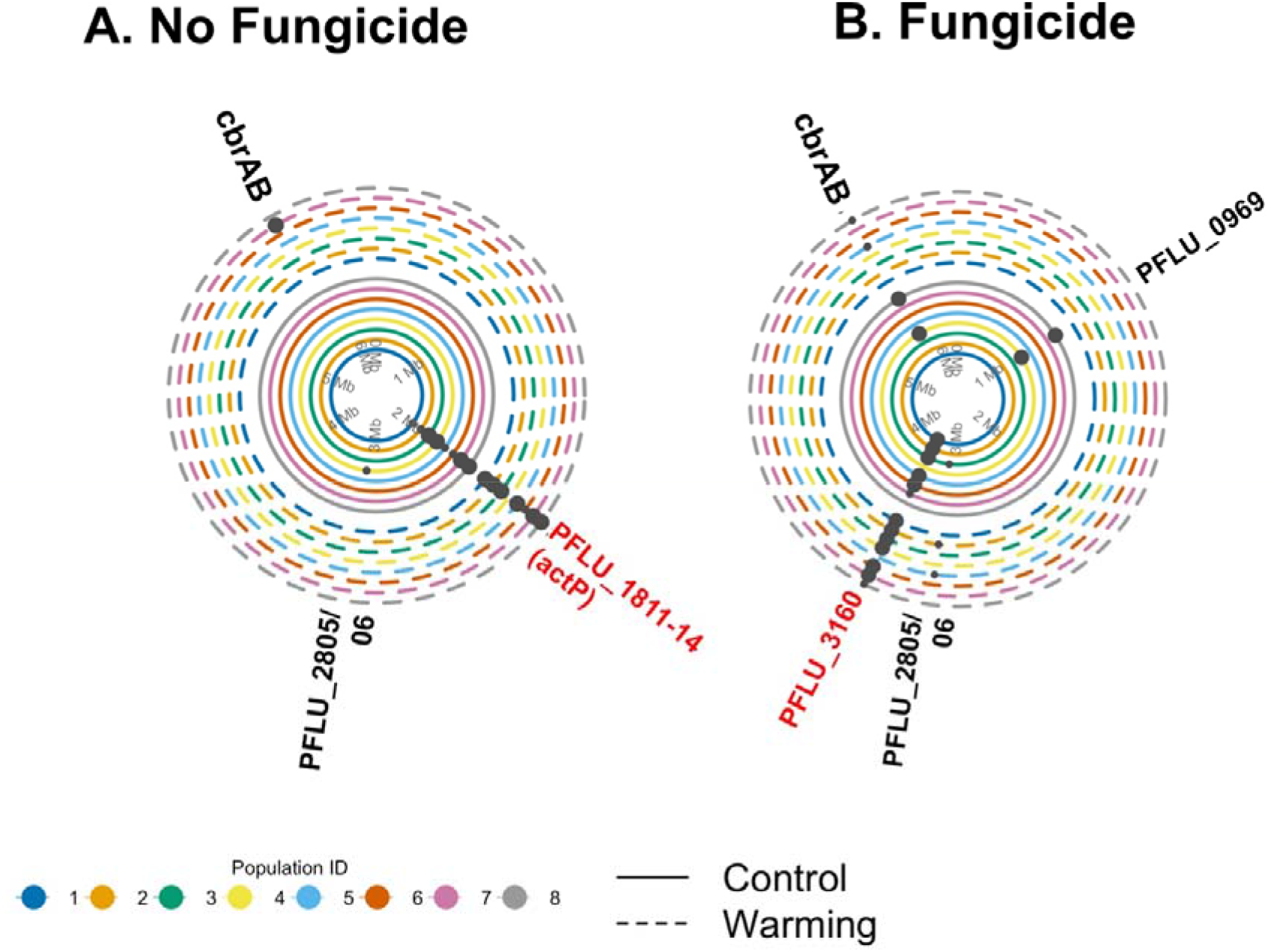
Mutations under positive selection as indicated by parallel genomic evolution in *P. fluorescens* populations evolved in fungicide-free (A) or (B) fungicide-treated soil. Each concentric circle corresponds to a replicate population in either the control (without fungicde) or treatment group (with fungicide), as indicated. Warmed and unwarmed populations are shown as broken lines and solid lines, respectively. Positions around each concentric circle, starting at the 12 o’clock position and in a clockwise direction, correspond to positions around the published *P. fluorescens* SBW25 single circular chromosome. Black points indicate the position of mutations; only those mutated in parallel (i.e. in >1 replicate population) are shown here. Point size corresponds to the number of clones harbouring the specific mutation per population (i.e. one or both sequenced clones). Mutation labels colured red are significantly associated with fungicide treatment: (*actP*, unique to fungicide-free populations) and *PFLU_3160* (unique to fungicide-treated populations). Mutations in *PFLU_1813, PFLU_1814 (actP)* and a large deletion in *PFLU_1811-actP* are grouped for clarity. A full list of mutations is available in supplementary data (S1 and S2).

Treatment-specific parallel evolution can suggest natural selection acting upon a specific gene or genes [41]. From our dataset of 5 parallel targets, two targets were significantly associated with fungicide treatment: i) *PFLU_3160*, unique to fungicide-treated populations and ii) *actP* – occurring only in non-fungicide treatments (Fisher’s exact test, bonferroni-adjusted: p<0.0001 for both targets, Figure 1. Mutations in *PFLU_3160* were identified in 24/32 clones across 13/16 fungicide-evolved populations and were detected as early as week 4 (fungicide+warming populations 1+5). *PFLU_3160* is a *P. aeruginosa mexS* ortholog which can regulate the expression of efflux pumps and porins [30, 31]. In contrast, mutations in *actP* or adjacent *PFLU_1813* were unique to fungicide-free populations, appearing in 23/30 clones across 15/16 populations. Mutations in *actP* have been previously shown to be advantageous in this soil environment [22], yet our findings suggest *actP* mutations are constrained in the presence of fungicide. Warming had no effect on the distribution of each of the 5 parallel targets across treatments (Fisher’s exact test, p>0.05 in all cases, Figure 1).

### 3.3 Selection for increased fungicide-resistance

Fungicide MIC was significantly affected by evolution treatment (ANOVA: F= 18.43, p < 0.0001, Figure 2a). Fungicide-evolved clones had increased fungicide MICs compared to fungicide-free controls, irrespective of whether fungicide was applied alone, or alongside warming (TukeyHSD; fungicide versus control: p < 0.001; fungicide + warming versus control: p < 0.0001, Figure 2a). Fungicide adaptation was not constrained by warming; there was no difference in MIC between fungicide populations evolved under control versus warming conditions (TukeyHSD; fungicide versus fungicide+warming: p > 0.5). For all tested clones, MIC’s ranged from 2 – 32 µg ml^−1^. Ancestral and control-evolved clones had an MIC of 2-8 µg ml^−1^, while fungicide-evolved clones had an MIC ranging from 4 -32 µg ml^−1^ (Table S1).

**Figure 2.**
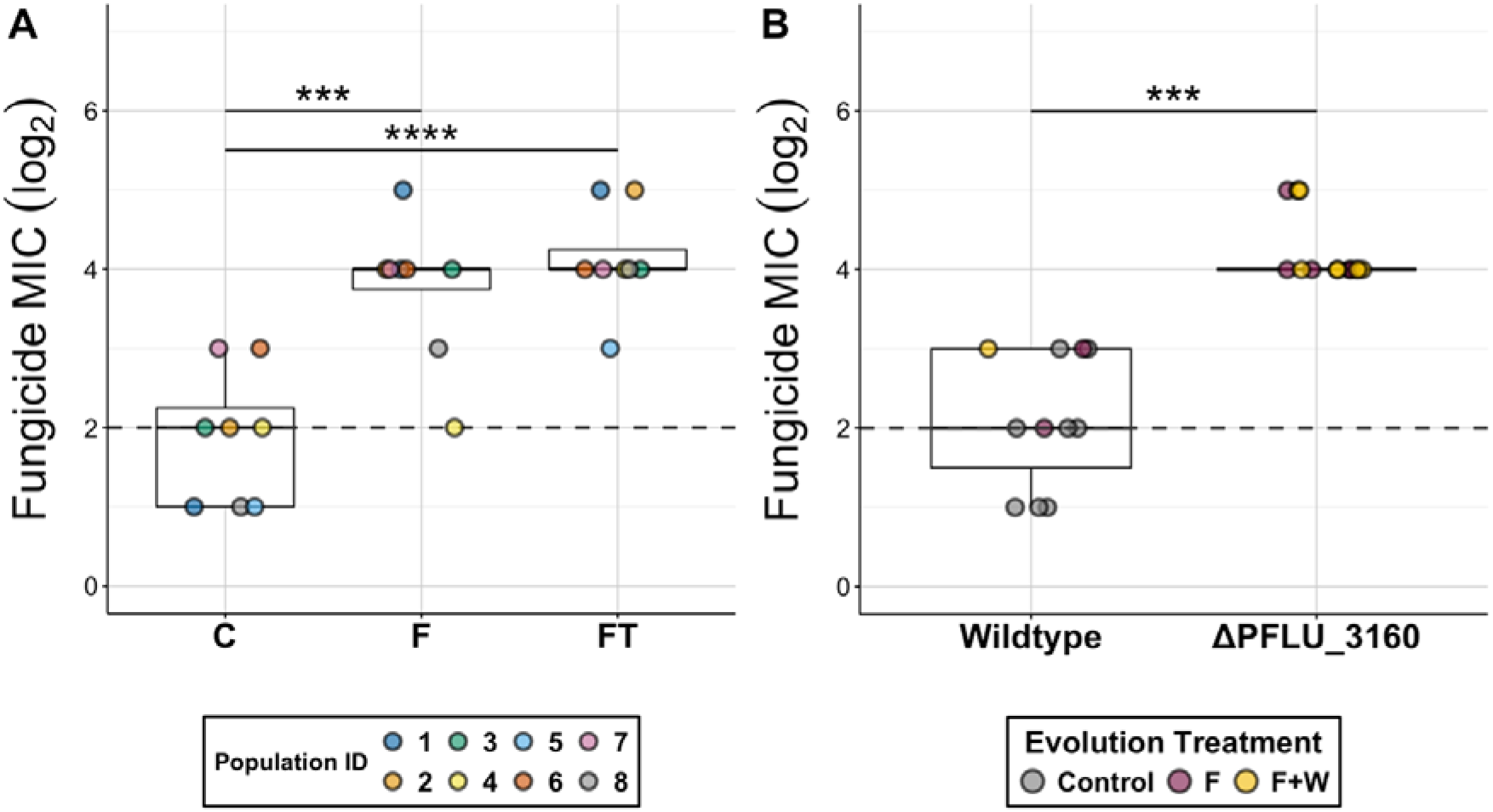
Fugicide resistance in evolved isolates. A) Log_2_ fungicide MIC for isolated *P. fluorescens* clones evolved under control, fungicide (F) or fungicide+warming (FT) treatments. Points are coloured according to population ID. B) The same dataset grouped by *PFLU_3160* mutation (ancestral or mutated). Points are coloured by evolutionary treatment. Individual points represent mode MIC for 3 technical replicates per clone. Dashed horizontal line shows mode ancestral Log_2_MIC value. Boxplots show the median (central line), interquartile range (box), and 1.5× interquartile range (whiskers). Lines connecting groups indicate significant differences between groups. Significance levels are denoted as: p < 0.05 (*), p < 0.01 (**), p < 0.001 (***), p < 0.0001 (****).

Our sequencing results suggested that mutations in *PFLU_3160* may be associated with increased resistance to fungicide. We compared fungicide MIC specifically for clones harbouring a *PFLU_3160* mutation versus clones with wildtype *PFLU_3160* (Figure 2b, Table S1). Our dataset comprised of fungicide-evolved clones with *PFLU_3160* mutation (n=13), ii) fungicide-evolved clones with wildtype *PFLU_3160* (n=3), iii) control (no-fungicide) evolved clones with wildtype *PFLU_3160* (n=8), and iv) ancestral clones of each population with wildtype *PFLU_3160* (n=8). *PFLU_3160* mutants had significantly increased MIC compared to wildtype overall (T-test, t=7.68, p<0.0001, Figure 2b, Table S1). Moreover, when both PFLU*_3160* status and treatment were included in the same linear model, *PFLU_3160* status was a better predictor of fungicide MIC than treatment (LM Effect of *PFLU_3160* status F_1,20_=14.339, p=0.001; NS effect of treatment F_2,22_=1.94, p>0.05). Across all treatments, fungicide MIC of wildtype *PFLU_3160* clones ranged between 2-8 fµg/ml, while clones harbouring a *PFLU_3160* mutation had increased MIC to 16-32 µg/ml (Table S1). Notably, the three fungicide-evolved clones with MIC similar to ancestor (4-8 µg/ml) were also the only three fungicide evolved clones with wildtype *PFLU_3160* (Table S2). Together, this data supports the hypothesis that *PFLU_3160* mutations enhance fungicide resistance in our experimental populations.

### 3.4 Fungicide selects for increased antimicrobial resistance

*PFLU_3160* is a *mexS* ortholog, and loss-of-function mutations in *mexS* have been implicated in conferring antimicrobial resistance in *Pseudomonas aeruginosa* [32, 33]. To test whether fungicide-selected mutations induce cross-resistance to antimicrobials, we performed antibiotic disk diffusion assays on the same sequenced clones subjected to fungicide resistance assays (section 3.3). Treatment-specific differences in reststance were observed for chloramphenicol (Kruskal-Wallis: χ^2^_2_=14.3, p<0.01), sulphatriad (Kruskal-Wallis: χ^2^_2_= 13.7, p < 0.01), nalidixic acid (Kruskal-Wallis: χ^2^_2_= 24.7, p < 0.0001) and tetracycline (Kruskal-Wallis: χ^2^_2_= 12.5, p< 0.05) (Figure 3, Table S2). Across these 4 antibiotics, fungicide treatments (applied alone or in combination with warming) had increased resistance compared to controls (Dunn test; fungicide versus control: p<0.05, fungicide+warming versus control: p<0.05, for all four antibiotics). There was no difference in resistance between fungicide treatments evolved under warming versus control temperatures (Dunn test; fungicide versus fungicide+warming, p>0.05 in all cases, Figure 3, Table S2). Hence, while fungicide increased resistance to 4/6 tested antibiotics, this was unaffected by experimental warming. No treatment-specific differences in resistance were observed for colistin sulphate (Kruskal-Wallis: χ^2^_2_= 2.3, p>0.05) or streptomycin (Kruskal-Wallis: χ^2^_2_= 6.34, p >0.05) (Figure 3, Table S2).

**Figure 3:**
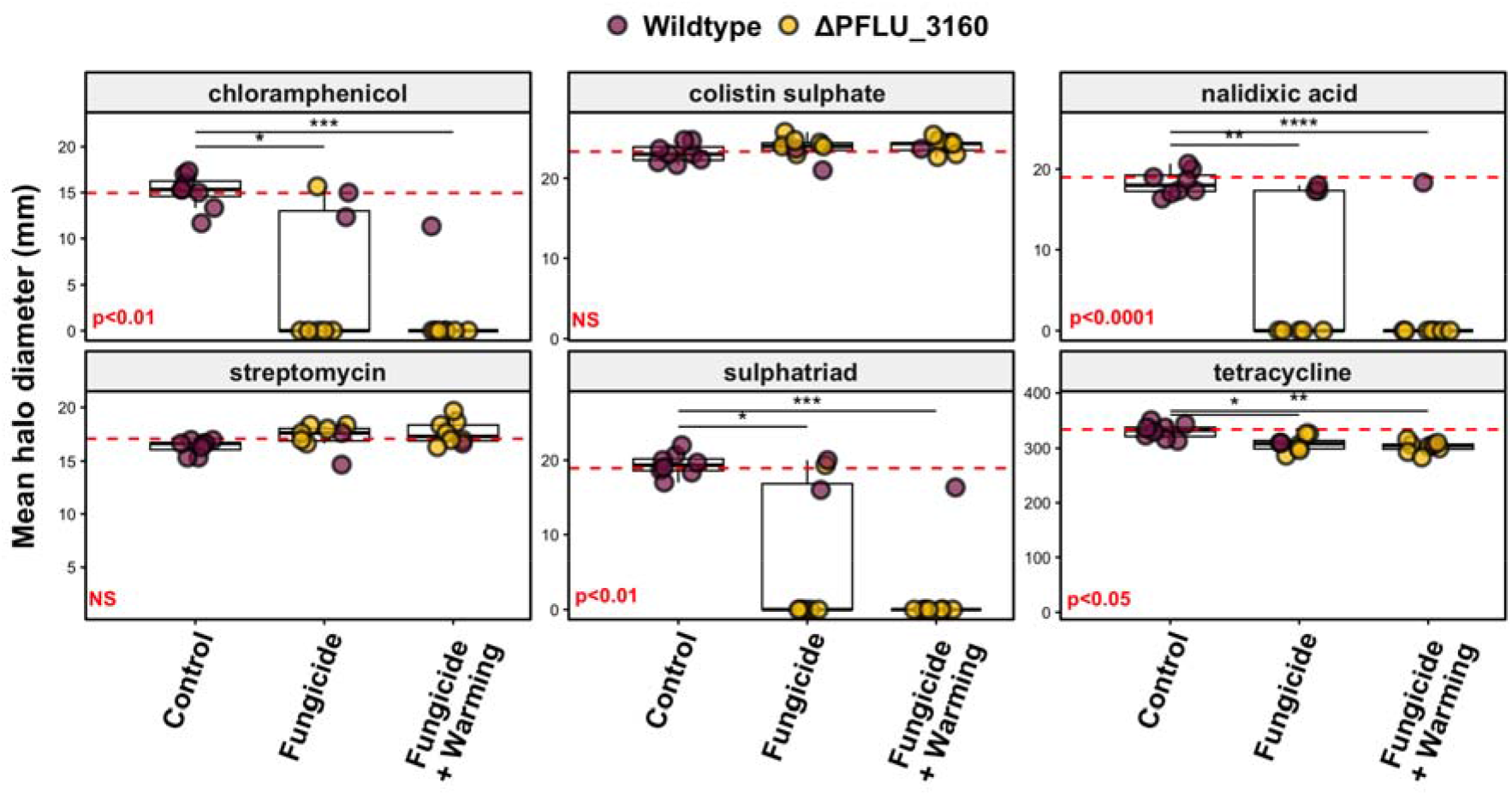
Mean halo diameter (mm) for *P. fluorescens* clones evolved under control, fungicide or fungicide+warming treatments, across six antimicrobials. Individual points represent mean halo diameter for 3 technical replicates per clone. Points are coloured according to whether that clone carries wildtype *PFLU_3160* or mutated *PFLU_3160*. Boxplots show the median (central line), interquartile range (box), and 1.5× interquartile range (whiskers). Lines connecting groups indicate significant differences based on Dunn posthoc tests. Overall Kruskal-Wallis P-values for each antibiotic are given in red. Significance levels are denoted as follows: p < 0.05 (*), p < 0.01 (**), p < 0.001 (***), p < 0.0001 (****). Red horizontal line shows mean value for 8 ancestral clones.

We next directly compared antimicrobial susceptibility of clones harbouring a *PFLU_3160* mutation versus control evolved clones with wildtype *PFLU_3160*. For chloramphenicol (Wilcox test; W=135, p<0.001), sulphatriad (Wilcox test; W=137, p<0.001), nalidixic acid (Wilcox test; W=555, p<0.0001) and tetracycline (Wilcox test; W=124, p<0.05) *PFLU_3160* mutants had significantly increased resistance versus wildtype (Figure 4, Table S2). Eleven out of 12 clones with mutations in *PFLU_3160* displayed complete resistance to chloramphenicol, nalidixic acid, and sulphatriad (i.e. zone of inhibition =0), while all wildtype clones remained susceptible to these antibiotics. The one exception to this finding was clone B3X (the only clone with a mutation in *PFLU_3161*, a *mexT* ortholog), which harboured a mutation in *PFLU_3160*, yet remained susceptible to chloramphenicol, nalidixic acid and sulphatriad (Table S2). Conversely, for streptomycin (Wilcox test; W=19.5, p<0.05), *PFLU_3160* mutants were more susceptible than wildtype clones, however this difference was very small (range of 1-2mm) (Figure 4), so may not be clinically relevant. Together, our data show fungicides can select for multidrug resistance, likely via selection for mutations in *PFLU_3160*.

**Figure 4:**
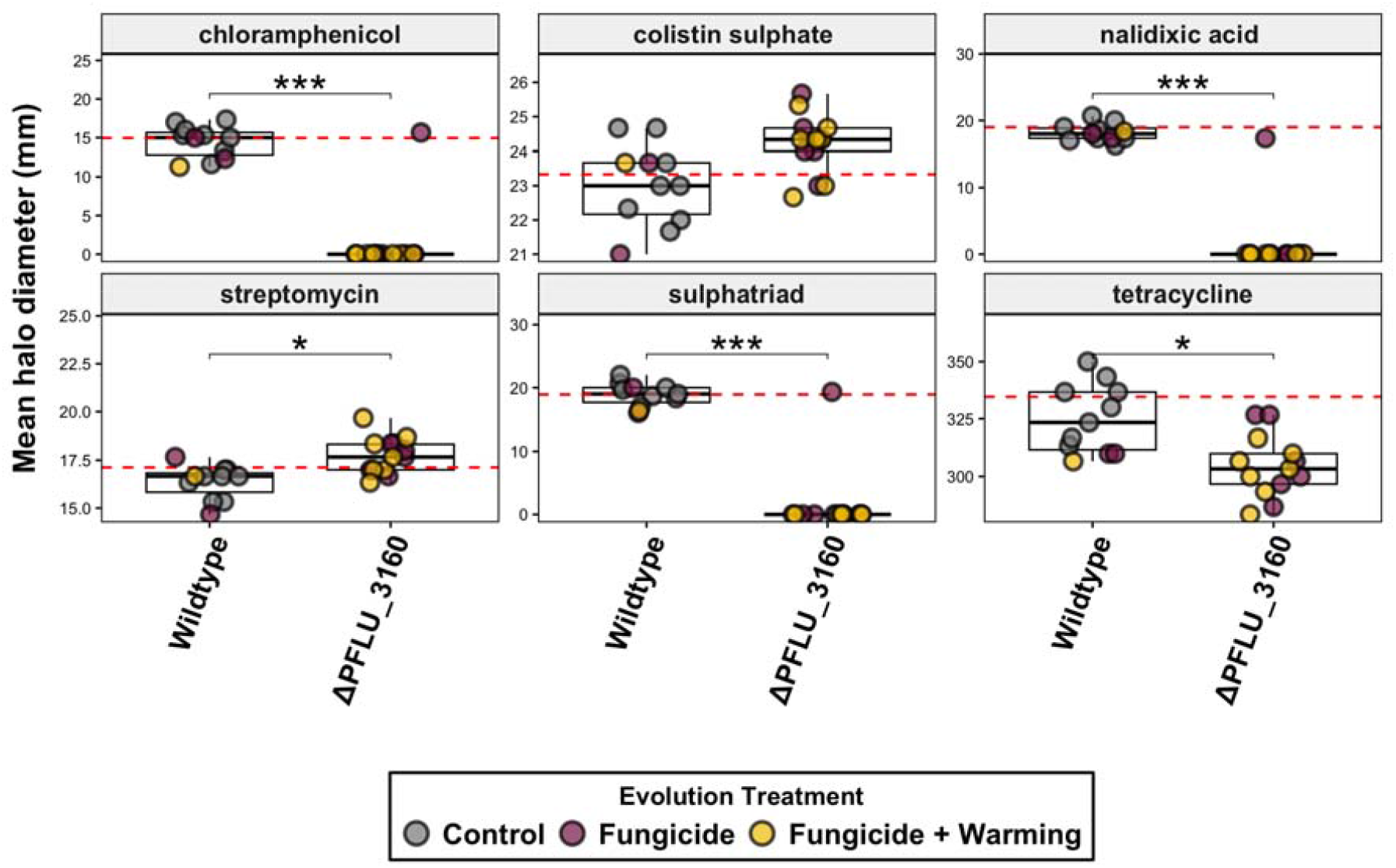
Antimicrobial susceptibility data from figure 3, grouped by *PFLU_3160* mutation status (wildtype v mutated). For chloramphenicol, sulphatriad, nalidixic acid and tetracycline (Wilcox test; W=124, p<0.05) *PFLU_3160* mutants had significantly increased resistance versus wildtype. 11/12 Clones with mutations in *PFLU_3160* displayed complete resistance to chloramphenicol, nalidixic acid, and sulphatriad (i.e. zone of inhibition =0), while all wildtype clones remained susceptible to these antibiotics. Conversely, for streptomycin, *PFLU_3160* mutants are more susceptible than wildtype clones, however this difference is very small (range of 1-2mm).Together, our data show fungicides can select for multidrug resistance, likely via selection for mutations in *PFLU_3160*.

## 4. Discussion

We investigated the impact of warming temperatures and fungicide accumulation on the growth and evolutionary trajectory of the PGPR *P. fluorescens*. Our data suggest that agricultural fungicides negatively impact bacterial growth, driving *de novo* mutations that confer increased resistance to fungicide, and inadvertently, antibiotics. Fungicide exposure drove a clear increase in fungicide resistance, with MICs rising from 2–8 µg ml^−1^ in ancestral and control-evolved clones to 8–32 µg ml^−1^ in fungicide-evolved populations. Strikingly, despite no prior antibiotic exposure, fungicide selection generated complete resistance to chloramphenicol and nalidixic acid, and frequently to sulphatriad, even at the high concentrations used in our assays. Our observations align with a growing body of evidence that pesticides can co-select for antibiotic resistance. For example, Xing et al. [34] showed that environmental pesticide exposure accelerates evolution toward higher antibiotic resistance in *E. coli*, while other studies demonstrate herbicide- and fungicide-mediated cross-selection of antibiotic resistance genes in microbial communities [35, 36]. Although horizontal gene transfer can contribute to such patterns in complex communities [35] our single-species design indicates that de novo mutation alone is sufficient to generate multidrug resistance. Together with work highlighting efflux-mediated cross-resistance between biocides and antibiotics [14, 37] our results reinforce the view that non-antibiotic agrochemicals can directly drive clinically relevant resistance phenotypes.

Warming did not alter selection for fungicide resistance: fungicide MICs and correlated antibiotic resistance did not differ between fungicide treatments evolved with or without warming temperatures. However, dual-stressor populations went extinct significantly earlier than single-stressor populations, despite harbouring clones that increased fungicide resistance. Thus, while warming did not constrain resistance evolution per se, it appears that the demographic benefit of resistance was not suffient for evolutionary rescue in dual-stressor populations. This decoupling of adaptive mutation from population persistence highlights an important principle: genotype-level benefits do not necessarily translate into population-level survival under multistressor conditions.

We found evidence for strong selection on the *mexS* ortholog *PFLU_3160. mexS* represses *mexT*, inhibiting the production of the MexEF-OprN multidrug efflux system [30, 33, 38] and increasing the expression of OprD porins [39]. In *P. aeruginosa*, overexpression of MexEF-OprN efflux pumps and reduced expression of OprD porins causes increased resistance to antibiotics through a mixture of efflux and reduced membrane permeability [30, 32, 33, 39-41]. However *mexS* mutations have not before been linked to fungicide resistance. Interestingly, the clone B3X was the only *PFLU_3160* mutant with increased fungicide resistance without a corresponding increase in antibiotic resistance. B3X was also the only clone with a mutation (SNP) in *PFLU_3161*, a *mexT* ortholog. This SNP mutation may have altered MexT structure, preventing the increased expression of MexEF-OprN while still reducing OprD porin expression, putatively explaining the observed phenotype of increased fungicide resistance (via OprD) but not antibiotic resistance (via MexEF-OprN).

Mutations in *actP*, an acetate-scavenging transporter [42] were widespread in fungicide-free control populations but were never detected in fungicide-free populations. We identied SNPs, insertion and large deletions in *actP* or *actP*-associated regions, consistent with loss-of-function. A previous study using the same *P. fluorescens* strain and experimental compost system, showed that *actP* mutations are beneficial in soil *per se*, yet are similarly constrained by the presence of a competing species, *P. putida [20]*. Why *actP* remains intact in the presence of fungicide (this study) or a competitor [20], remains unclear. In *Neisseria meningitidis*, acetate can protect against oxidative stress [43], suggesting that in *P. fluorescens*, acetate uptake via ActP might help buffer cells against reactive oxygen species generated by mancozeb (a component of Fubol Gold) [44] or by metabolic stress arising from nutrient competition [44]. One caveat is that we sequenced clones from the last viable timepoint of our evolution experiment, which was earlier for fungicide-evolved populations. Since fungicide-free clones had a longer period of evolution (and on average, higher population densities), it is possible that *actP* mutations would have eventually evolved in fungicide-treated populations. However, given that Hall et al. [20] found similarly strong selection for *actP* mutations in the absence, but not presence of a (biotic) stressor, and the clear environment-dependent benefits of *actP* mutations quantified by [22], there is good support for the general role of environmental stress-dependent fitness effects of *actP*.

Our study, alongside a growing body of work [11, 36], challenges the prevailing paradigm that antibiotics are the sole or even dominant drivers of AMR. It is now clear that selection for AMR is shaped by a complex mixture of environmental stressors, including pesticides, biocides, heavy metals, microplastics, and other anthropogenic contaminants [11]. Our findings show that fungicide exposure alone, in the complete absence of antibiotics, can select de novo mutations generating multidrug resistance phenotypes. This suggests that reducing antibiotic use, while essential, may be insufficient to curb AMR if co-selective agents in agricultural environments remain unaddressed.

