## Supplementary material for "Fungicide drives *de novo* evolution of multidrug resistance in the plant growth promoting rhizobacterium, *Pseudomonas fluorescens*": Figure S

**Supplementary Figures and Tables**

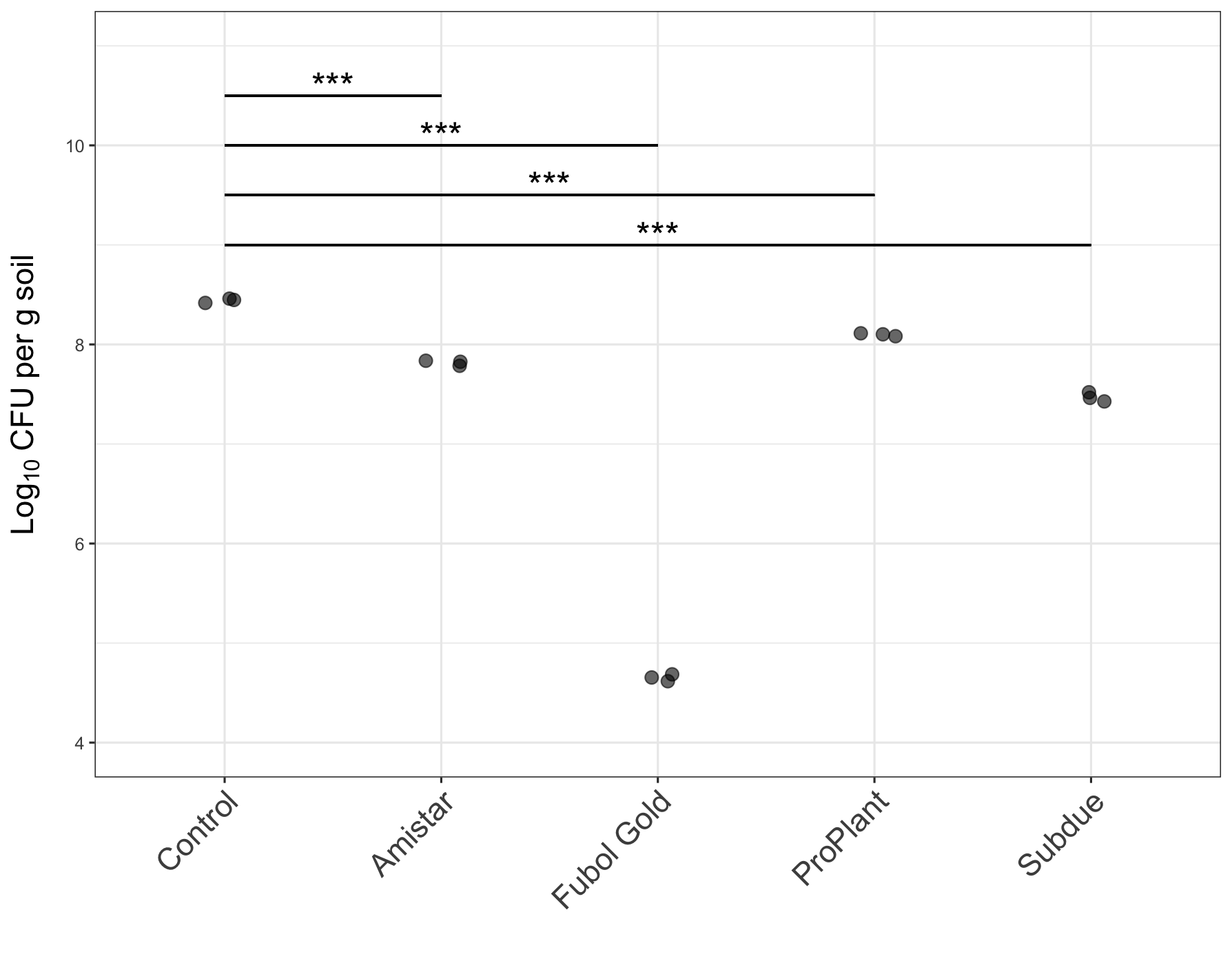

**Supplementary figure S1**: The effect of four different fungicides on final densities of *P. fluorescens* in soil microcosms (Log_10_ CFU g^-1^). All fungicide formulations significantly decreased *P. fluorescens* densities compared to fungicide-free controls (see main text for statistics). Assays were replicated in triplicate. Asterisks denote significant pairwise t-test comparisons, with Bonferroni adjustment .

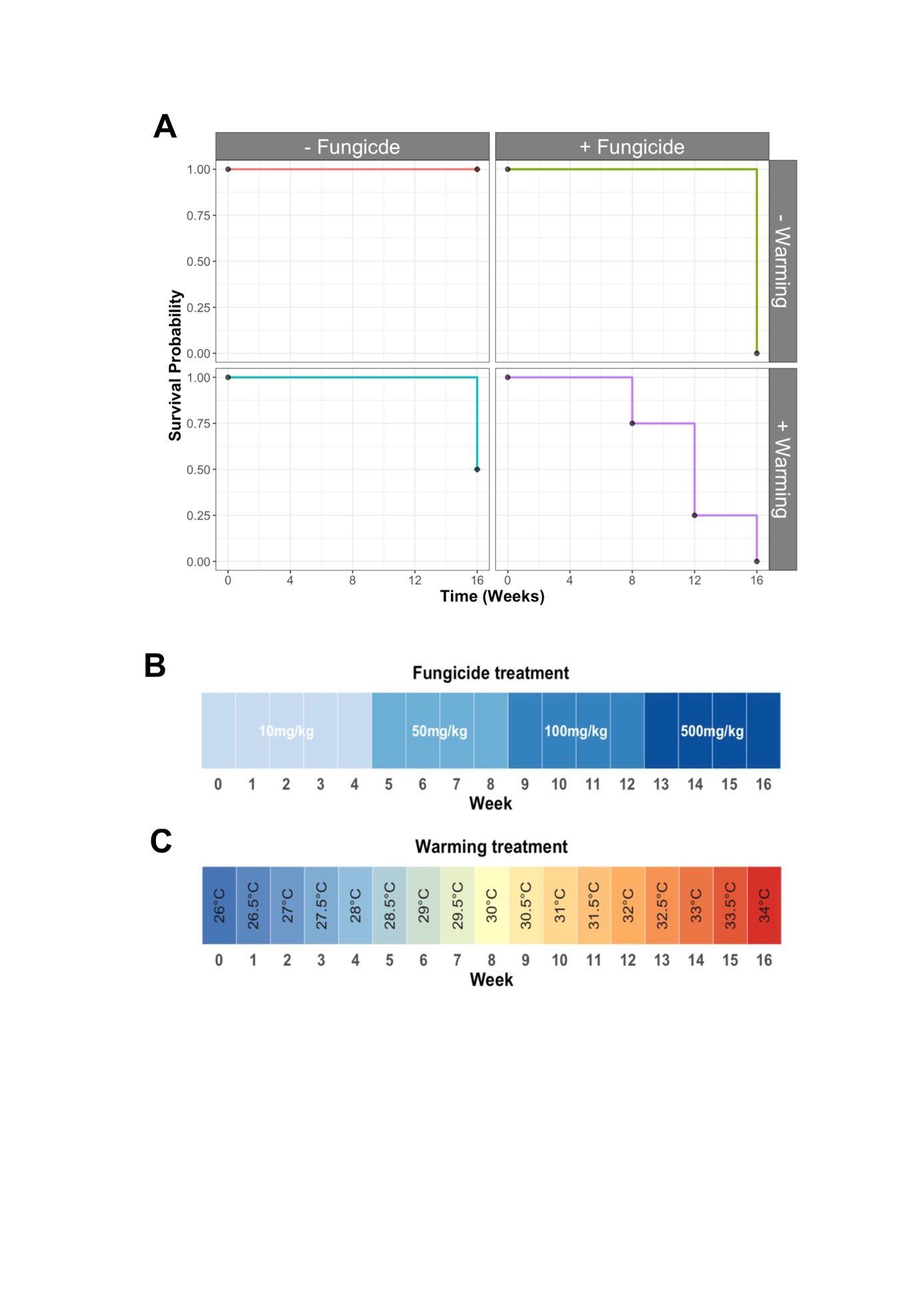

**Supplementary figure S2**: **A)** Kaplan–Meier survival curves showing the survival probability of experimental populations over time under four treatments: control (no warming, no fungicide), fungicide only, warming only, and dual stress (warming + fungicide). Dual-stress populations went extinct earliest and most completely, whereas single-stressor populations experienced intermediate losses and all control populations remained viable. See main text for statistics. **B)** Fungicide concentrations in soil increased every 4 weeks, to a maximum of 500 mg kg^-1^ at weeks 12-16. **C)** In warming treatments, temperatures increased by 0.5°C weekly until a maximum temperature of 34°C by week 16.

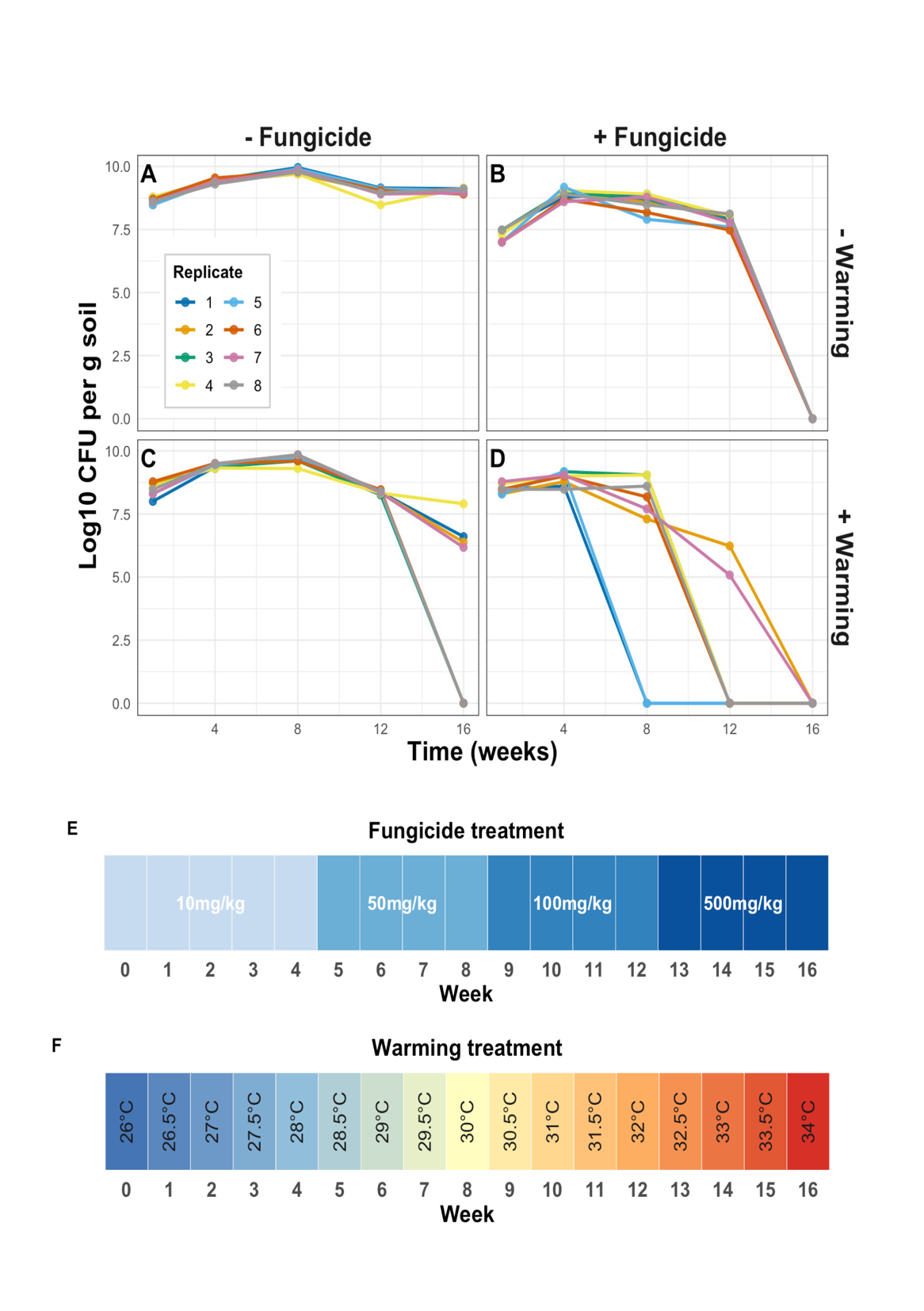

**Supplementary figure S3**: Population densities (Log_10_ CFU g^-1^ soil) over the course of a 16 week evolution experiment. Populations were evolved under control conditions (A), with fungicide (B), warming (C) or both fungicide and warming (D). Individual lines show densities for a single evolving population in soil. Fungicide concentrations in soil increased every 4 weeks, to a maximum of 500 mg kg^-1^ at weeks 12-16 (E). In warming treatments, temperatures increased by 0.5°C weekly until a maximum temperature of 34°C by week 16 (F). Dual-stressor populations showed the greatest decline in population densities, with extinctions as early as week 8.

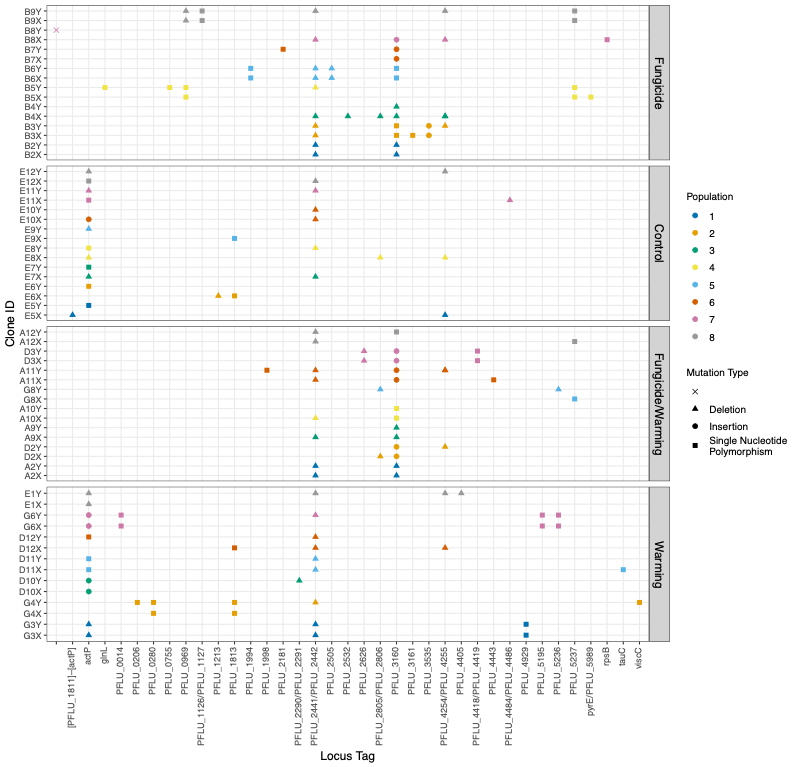

**Supplementary figure S4**: To identify treatment-specific patterns of mutation, we sequenced the genome of two clones from each evolved population (Clone ID; y-axis). Locus tag/gene name is shown on x-axis. Clones are separated by treatment in which they evolved (Fungicide, Warming, Fungicide+Warming, Control). Across evolved clones, we detected 152 mutations comprising gene deletions (triangles), insertions (circles) and single nucleotide polymorphisms (squares). Evolved clones had between 0 and 6 mutations with a median of 2 mutations per clone. We did not detect any mutations in one clone (B8Y; fungicide-only treatment). Two targets consisted of large deletions in repeat-rich intergenic regions, which are prone to spurious mutation calls and should be interpreted with caution (*PFLU_2441-PFLU_2442* and *PFLU_4254- PFLU_4255).* Colour corresponds to a particular replicate population. Further information on specific mutations is available in Supplementary data S1.

**Supplementary table** **1:** Mode fungicide minimum inhibitory concentrations (MIC) for evolved and all 8 ancestral *P. fluorescens* clones. MIC’s are denoted via an orb colour, with green being the most resistant and red being the least resistant to fungicide, respectively. All clones were subjected to whole genome resequencing and details on *PFLU_3160* (*mexS* ortholog) mutation status is given. All clones harbouring a PFLU_3160 mutation have increased resistance to fungicide. Mutation type: DEL = Deletion; SNP = Single Nucleotide Polymorphism; INS = Insertion; - = no mutation.

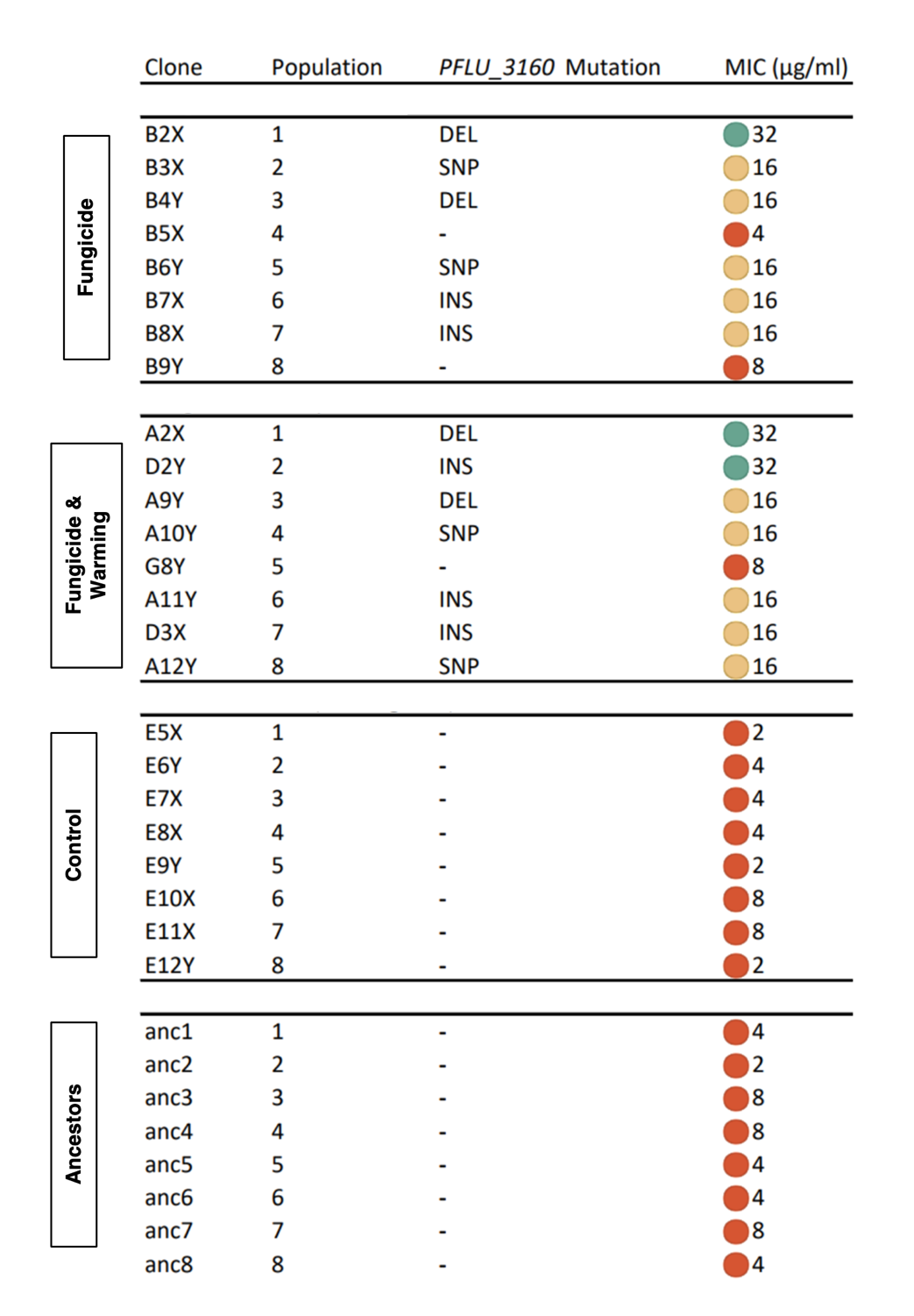

**Supplementary table S2:** Antimicrobial zone of inhibition in mm (ZOI) for evolved and ancestral *P. fluorescens* clones. Resistance level is denoted via an orb colour, with green being the most susceptible and red being completely resistant, respectively. harbouring mutations in *PFLU_3160* are associated with complete resistance (ZOI = 0mm) to chloramphenicol, sulphatriad and nalidixic acid. The following disc concentrations were used: Chloramphenicol (30 µg), Sulphatriad (200 µg), Tetracycline (25 µg), Nalidixic acid (30 µg). Mutation type: DEL = Deletion; SNP = Single Nucleotide Polymorphism; INS = Insertion; - = no mutation. All ancestral clones were within the same range as control clones (See Figure 3 in main text for statistics).

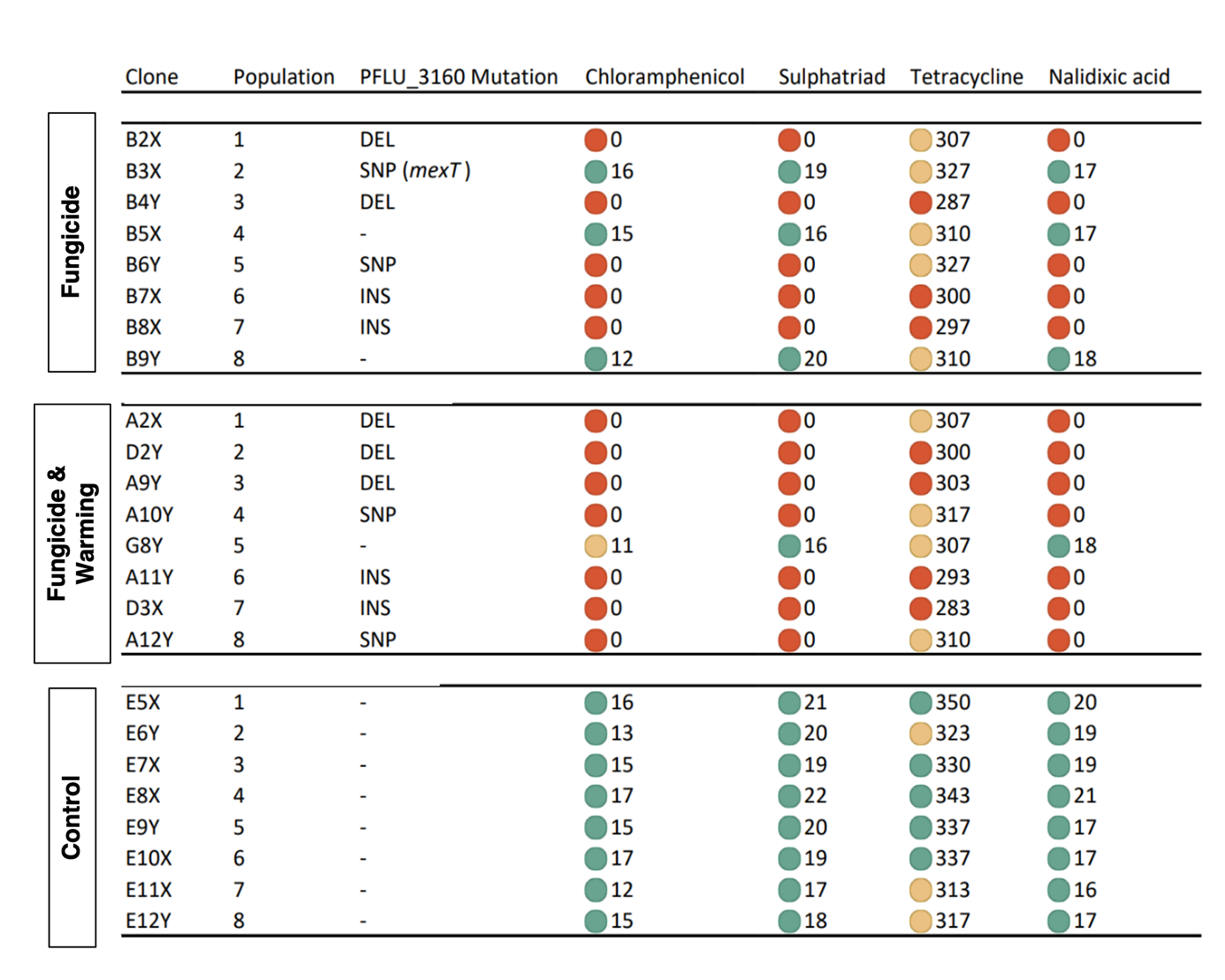
