## Supplementary Methods for "Fungicide drives *de novo* evolution of multidrug resistance in the plant growth promoting rhizobacterium, *Pseudomonas fluorescens*"

PCR identification of isolated clones: To ensure soil-isolated clones were our inoculated *P. fluorescens* strain. we isolated 12 clones at random (3 from individual populations of all four treatments) for PCR analysis of the mini-Tn7 Gent^R^ cassette (a unique sequence unlikely to be found in native microbes). The gentamicin resistance transposon was amplified using a master mix of the primers Tn7-GlmS-pf1F ('CGTCATCAACATGCCGCACATC') and Tn7R109 ('CAGCATAACTGGACTGATTTCAG') (44). Colony PCR was performed using GoTaq G2 Green Master Mix. A 48 h LBA grown culture was touched with a sterile pipette tip and mixed into 25 µl of the master mix (along with a +/- control) and run on a thermal cycler with the following program: initial denaturation 95 °C for 5 min, followed by thirty cycles of 95°C for 30 sec, 58°C for 30 sec, 72°C for 1 minute, followed by a final elongation at 72°C for 5 min. The PCR product was run on a 1% agarose (Bioline) and TAE (Sigma) gel with a 100 bp ladder (New England Biolabs) at 100 volts before imaging under blue light. All tested soil clones showed a band at 300 bp, indicative of the mini-Tn7 construct, suggesting that the focal strain had been successfully recovered.
